# Multi-tissue probabilistic fine-mapping of transcriptome-wide association study identifies cis-regulated genes for miserableness

**DOI:** 10.1101/682633

**Authors:** Calwing Liao, Veikko Vuokila, Alexandre D Laporte, Dan Spiegelman, Patrick A. Dion, Guy A. Rouleau

## Abstract

Miserableness is a behavioural trait that is characterized by strong negative feelings in an individual. Although environmental factors tend to invoke miserableness, it is common to feel miserable ‘for no reason’, suggesting an innate, potential genetic component. Currently, little is known about the functional relevance of common variants associated with miserableness. To further characterize the trait, we conducted a transcriptome-wide association study (TWAS) on 373,733 individuals and identified 104 signals across brain tissue panels with 37 unique genes. Subsequent probabilistic fine-mapping prioritized 95 genes into 90%-credible sets. Amongst these prioritized hits, *C7orf50* had the highest posterior inclusion probability of 0.869 in the brain cortex. Furthermore, we demonstrate that many GWAS hits for miserableness are driven by expression. To conclude, we successfully identified several genes implicated in miserableness and highlighted the power of TWAS to prioritize genes associated with a trait.

**Short summary:** The first transcriptome-wide association study of miserableness identifies many
genes including *c7orf50* implicated in the trait.

## Introduction

Miserableness is characterized by emotional distress, typically caused by an event invoking negative feelings in an individual^1^. Although environmental factors may cause feelings of miserableness, it is common for certain individuals to feel miserable independent of environmental influences, suggesting that underlying biologic and genetic factors might play a role in miserableness. Several large-scale genome-wide association studies (GWAS) have focused on characterizing the genetics of neuroticism items such as miserableness^2,3^. No genetic factors fully overlap both miserableness and other neuroticism traits, which suggests that there are variants and/or genes that are specific to miserableness^4^. For instance, previous interrogation of neuroticism items found that miserableness had the highest genetic correlation with feeling “fed-up” (0.88) and “experiencing mood swings” (0.86), suggesting that there are genetic differences^4^. However, identifying the biological functionality of these variants remains difficult.

Transcriptomic imputation has recently been used to integrate genotype and expression data to identify predictive cis-eQTLs, which in turn can be applied to independent datasets in a tissue-specific manner^5^. Transcriptome-wide association studies (TWAS) can identify cis-eQTL regulated genes by integrating expression panels from large consortia such as GTEx and the CommonMind Consortium (CMC) with GWAS data^6^. Furthermore, probabilistic fine-mapping of these TWAS hits can be done by modelling correlation among TWAS hits and assigning a probability for each gene within the risk region to prioritize genes^7^.

To understand cis-eQTL regulated candidate genes in miserableness, we conducted a TWAS consisting of 373,733 individuals using the summary statistics from a recent study dissecting the genetic heterogeneity of neuroticism^4^. The participants were asked whether they tend to ‘feel miserable’ for no particular reason. A total of 104 TWAS signals with 37 unique genes were transcriptome-wide significant across brain tissue panels. Probabilistic fine-mapping identified a number of putatively causal genes, including *C7orf50* in *GPX1* in the frontal cortex with a posterior inclusion probability (PIP) of 0.869 and *MTCH2* in the nucleus accumbens with a PIP of 0.573. Conditioning on the transcriptome-wide significant hits showed that the hits accounted for >75% of the GWAS signal present. To conclude, miserableness genetics is explained partially by cis-eQTLs driving the expression of certain genes. Additional studies are needed to characterize other sources of non-coding variation such as 3’-aQTLs.

## Results

### Transcriptome-wide association study identifies 104 signals associated with miserableness

A multi-tissue TWAS using FUSION identified 104 TWAS signals associated with miserableness after Bonferroni-correction (Table 1, Figure 1). Across all signals, there were 37 genes identified in at least one imputation panel (Supplementary table 1). Amongst all signals, the top five hits in chronological order consisted of *GPX1* (P_Bonferroni_= 5.18 × 10^-10^) in the frontal cortex, *RNF123* (P_Bonferroni_=3.67 × 10^-8^) in the cerebellum, *GPX1* (P_Bonferroni_=3.09 × 10^-7^) in the cerebellum, *RBM6* (P=5.12×10^-7^) in the caudate basal ganglia, and *MST1R* (P_Bonferroni_=1.29 × 10^-6^) in the prefrontal cortex. Amongst the signals, several implicated RNA-coding genes, reinforcing the notion that non-coding genes are relevant to complex traits (Table 1).

**Table 1.** Multi-tissue significant TWAS hits for miserableness.

| Gene | Tissue | eQTL | Z-score | Uncorrected P-value | Bonferroni P-value |
| --- | --- | --- | --- | --- | --- |
| AF131215.2 | ROSMAP Brain | rs17724226 | -5.57 | 2.51E-08 | 3.92E-03 |
|  | Cerebellar Hemisphere | rs17724226 | -5.93 | 3.11E-09 | 4.86E-04 |
| AF131215.3 | ROSMAP Brain | rs2409745 | -6.13 | 8.79E-10 | 1.37E-04 |
| AMT | Brain Cortex | rs3448 | -3.49 | 2.05E-07 | 3.20E-02 |
|  | Caudate Basal Ganglia | rs9834003 | -4.89 | 7.18E-08 | 1.12E-02 |
|  | ROSMAP Brain | rs7619016 | -11.57 | 6.45E-08 | 1.01E-02 |
|  | Hippocampus | rs17689471 | -4.07 | 1.89E-08 | 2.95E-03 |
| ARHGAP27 | Nucleus Accumbens Basal Ganglia | rs17689471 | 5.94 | 2.91E-09 | 4.54E-04 |
| C1QTNF4 | Brain Cortex | rs12286721 | 6.41 | 1.41E-10 | 2.20E-05 |
| CRHR1-IT1 | Brain Cortex | rs17689471 | 5.17 | 2.37E-07 | 3.70E-02 |
|  | Nucleus Accumbens Basal Ganglia | rs8072451 | 5.74 | 9.67E-09 | 1.51E-03 |
|  | Caudate Basal Ganglia | rs17763086 | 5.88 | 4.10E-09 | 6.40E-04 |
|  | Hypothalamus | rs17763086 | 5.99 | 2.06E-09 | 3.22E-04 |
|  | Frontal Cortex BA9 | rs17763086 | 6.02 | 1.71E-09 | 2.67E-04 |
|  | Putamen basal ganglia | rs17689471 | 6.05 | 1.43E-09 | 2.23E-04 |
|  | Hippocampus | rs17689471 | 6.43 | 1.31E-10 | 2.05E-05 |
| ERI1 | Dorsolateral prefrontal cortex | rs7826478 | 5.95 | 2.74E-09 | 4.28E-04 |
| FAM167A | Dorsolateral prefrontal cortex | rs7844834 | 5.44 | 5.47E-08 | 8.54E-03 |
|  | ROSMAP Brain | rs10102600 | 5.93 | 3.08E-09 | 4.81E-04 |
| FAM212A | Cerebellum | rs7372966 | 5.860692 | 4.61E-09 | 7.20E-04 |
| FAM66A | ROSMAP Brain | rs12676145 | -5.39 | 7.16E-08 | 1.12E-02 |
| FAM66D | ROSMAP Brain | rs12550733 | 5.21 | 1.90E-07 | 2.97E-02 |
| FAM85B | ROSMAP Brain | rs2945251 | 5.64 | 1.72E-08 | 2.69E-03 |
| FAM86B1 | Dorsolateral prefrontal cortex | rs11998678 | -5.46 | 4.90E-08 | 7.65E-03 |
| FAM86B3P | Frontal Cortex BA9 | rs2948286 | 5.5 | 3.69E-08 | 5.76E-03 |
|  | ROSMAP brain | rs2945230 | 5.65 | 1.57E-08 | 2.45E-03 |
|  | Cerebellar hemisphere | rs876954 | 6.25 | 4.14E-10 | 6.47E-05 |
|  | Cerebellum | rs2980436 | 6.5 | 7.99E-11 | 1.25E-05 |
|  | Brain Cortex | rs2945230 | 6.5 | 7.91E-11 | 1.24E-05 |
| FNBP4 | ROSMAP Brain | rs7929014 | -6.23 | 4.77E-10 | 7.45E-05 |
| FTSJ2 | Brain Cortex | rs3757440 | 5.805551 | 6.42E-09 | 1.00E-03 |
| GPX1 | Caudate Basal Ganglia | rs11130186 | 5.680812 | 1.34E-08 | 2.09E-03 |
|  | Cerebellar hemisphere | rs1873625 | 5.847834 | 4.98E-09 | 7.78E-04 |
|  | Dorsolateral prefrontal cortex | rs4955430 | 5.944679 | 2.77E-09 | 4.33E-04 |
|  | ROSMAP brain | rs4955430 | 6.35984 | 2.02E-10 | 3.15E-05 |
|  | Cerebellum | rs9853683 | 7.035615 | 1.98E-12 | 3.09E-07 |
|  | Frontal cortex BA9 | rs7617480 | 7.878335 | 3.32E-15 | 5.18E-10 |
| KANSL1-AS1 | Cerebellum | rs17689918 | 5.51 | 3.65E-08 | 5.70E-03 |
|  | Hippocampus | rs8072451 | 5.84 | 5.23E-09 | 8.17E-04 |
|  | Brain Cortex | rs17689918 | 5.86 | 4.53E-09 | 7.07E-04 |
|  | Nucleus accumbens basal ganglia | rs8072451 | 5.87 | 4.40E-09 | 6.87E-04 |
|  | Frontal cortex BA9 | rs8072451 | 5.87 | 4.35E-09 | 6.79E-04 |
|  | Putamen basal ganglia | rs8072451 | 5.88 | 4.21E-09 | 6.57E-04 |
|  | Caudate basal ganglia | rs17689471 | 5.88 | 4.04E-09 | 6.31E-04 |
|  | Cerebellar hemisphere | rs17689918 | 6.01 | 1.83E-09 | 2.86E-04 |
|  | Hypothalamus | rs8072451 | 6.12 | 9.41E-10 | 1.47E-04 |
| LRP4 | Dorsolateral prefrontal cortex | rs10838635 | -5.59 | 2.31E-08 | 3.61E-03 |
| LRRC37A | Cerebellum | rs169201 | 5.45 | 5.15E-08 | 8.04E-03 |
| LRRC37A2 | Brain cortex | rs199534 | 5.43 | 5.61E-08 | 8.76E-03 |
|  | Cerebellum | rs169201 | 5.66 | 1.49E-08 | 2.33E-03 |
|  | Hippocampus | rs169201 | 5.67 | 1.41E-08 | 2.20E-03 |
|  | Cerebellar hemisphere | rs169201 | 5.7 | 1.22E-08 | 1.91E-03 |
|  | Frontal cortex BA9 | rs199439 | 5.78 | 7.63E-09 | 1.19E-03 |
|  | Nucleus accumbens basal ganglia | rs199439 | 5.79 | 6.90E-09 | 1.08E-03 |
|  | Putamen basal ganglia | rs199439 | 5.79 | 6.88E-09 | 1.07E-03 |
|  | Hypothalamus | rs199439 | 5.81 | 6.14E-09 | 9.59E-04 |
| LRRC37A4P | Brain cortex | rs17689918 | -5.38 | 7.49E-08 | 1.17E-02 |
|  | Caudate basal ganglia | rs8072451 | -5.86 | 4.68E-09 | 7.31E-04 |
|  | Putamen basal ganglia | rs8072451 | -5.86 | 4.68E-09 | 7.31E-04 |
|  | Hippocampus | rs8072451 | -5.86 | 4.56E-09 | 7.12E-04 |
|  | Nucleus accumbens basal ganglia | rs8072451 | -5.86 | 4.51E-09 | 7.04E-04 |
|  | Cerebellar hemisphere | rs8072451 | -5.87 | 4.45E-09 | 6.95E-04 |
|  | Cerebellum | rs17689471 | -5.87 | 4.44E-09 | 6.93E-04 |
|  | Frontal cortex BA9 | rs17763086 | -5.91 | 3.45E-09 | 5.39E-04 |
|  | Hypothalamus | rs8072451 | -6.01 | 1.83E-09 | 2.86E-04 |
| MAPT | Cerebellar hemisphere | rs17763086 | -5.88 | 4.10E-09 | 6.40E-04 |
|  | Frontal cortex BA9 | rs17689882 | -5.9 | 3.55E-09 | 5.54E-04 |
|  | Dorsolateral prefrontal cortex | rs17689471 | -6.24 | 4.40E-10 | 6.87E-05 |
|  | Brain cortex | rs17689882 | -6.36 | 2.04E-10 | 3.19E-05 |
| MST1R | Dorsolateral prefrontal cortex | rs2526388 | 5.226389 | 1.73E-07 | 2.70E-02 |
|  | Cerebellum | rs2856236 | 6.563222 | 5.27E-11 | 8.23E-06 |
|  | ROSMAP brain | rs11713193 | 6.833558 | 8.28E-12 | 1.29E-06 |
| MTCH2 | Brain cortex | rs7947730 | -5.15 | 2.55E-07 | 3.98E-02 |
|  | ROSMAP brain | rs4752856 | -5.55 | 2.85E-08 | 4.45E-03 |
| MTMR9 | ROSMAP brain | rs2246606 | 5.44 | 5.39E-08 | 8.42E-03 |
| NICN1 | Nucleus accumbens basal ganglia | rs11130186 | 5.260183 | 1.44E-07 | 2.25E-02 |
| PLEKHM1 | Brain Cortex | rs11012 | 5.21 | 1.94E-07 | 3.03E-02 |
|  | Cerebellar hemisphere | rs8072451 | -5.9 | 3.63E-09 | 5.67E-04 |
|  | Cerebellum | rs17689918 | -6.39 | 1.65E-10 | 2.58E-05 |
| RBM6 | ROSMAP brain | rs6765484 | -5.375061 | 7.66E-08 | 1.20E-02 |
|  | Dorsolateral prefrontal cortex | rs9311446 | -5.785099 | 7.25E-09 | 1.13E-03 |
|  | Putamen basal ganglia | rs2240329 | -5.843777 | 5.10E-09 | 7.96E-04 |
|  | Frontal cortex BA9 | rs2681780 | 5.852691 | 4.84E-09 | 7.56E-04 |
|  | Cerebellar hemisphere | rs11713193 | -5.870895 | 4.33E-09 | 6.76E-04 |
|  | Nucleus accumbens basal ganglia | rs4688755 | -5.912526 | 3.37E-09 | 5.26E-04 |
|  | Cerebellum | rs6446193 | -6.056184 | 1.39E-09 | 2.17E-04 |
|  | Brain cortex | rs6765484 | -6.185421 | 6.19E-10 | 9.67E-05 |
|  | Caudate basal ganglia | rs2014830 | -6.965004 | 3.28E-12 | 5.12E-07 |
| RNF123 | Dorsolateral prefrontal cortex | rs2352974 | 5.465525 | 4.62E-08 | 7.21E-03 |
|  | Cerebellar hemisphere | rs7634902 | 5.474334 | 4.39E-08 | 6.86E-03 |
|  | Brain cortex | rs7634902 | 5.559422 | 2.71E-08 | 4.23E-03 |
|  | Frontal cortex BA9 | rs7634902 | 5.924265 | 3.14E-09 | 4.90E-04 |
|  | ROSMAP brain | rs2352974 | 6.269043 | 3.63E-10 | 5.67E-05 |
|  | Caudate basal ganglia | rs11716575 | 6.449052 | 1.13E-10 | 1.76E-05 |
|  | Cerebellum | rs11716575 | 7.327323 | 2.35E-13 | 3.67E-08 |
| RP11-l65J3.6 | Cerebellum | rs7025805 | 5.16009 | 2.47E-07 | 3.86E-02 |
| RP11-481A20.11 | ROSMAP brain | rs4841662 | -5.51 | 3.63E-08 | 5.67E-03 |
| RP11-750H9.5 | Cerebellar hemisphere | rs896817 | -6.05 | 1.46E-09 | 2.28E-04 |
| SEMA3F | ROSMAP brain | rs2247510 | -5.798856 | 6.68E-09 | 1.04E-03 |
| SLC38A3 | ROSMAP brain | rs1061474 | -5.503226 | 3.73E-08 | 5.82E-03 |
| UBXN2A | Dorsolateral prefrontal cortex | rs12616678 | -5.354554 | 8.58E-08 | 1.34E-02 |
|  | ROSMAP brain | rs7588286 | -5.926861 | 3.09E-09 | 4.83E-04 |
| XKR6 | ROSMAP brain | rs7821914 | -5.52 | 3.41E-08 | 5.33E-03 |

### Top GWAS signals are largely explained by expression

To determine how much GWAS signal remains after the expression association from TWAS is removed, conditional testing was done for transcriptome-wide significant TWAS signals. For most of the GWAS signals, the expression accounted for >75% of the variance (Table 2, Supplementary Figures 1-14). For *RP11-74E22.6, RP11-798G7.5*, and *DNF-AS*, conditioning on the expression accounted for 100% of the signal. There were less TWAS signals compared to the GWAS significant signals for miserableness, suggesting that other genetic mechanisms are driving those signals.

**Table 2.** Variance explained of GWAS signals by conditioning on expression for miserableness.

| Lead GWAS SNP | Conditioned on | Tissue | Before conditioning P-value | After conditioning P-value | Variance explained |
| --- | --- | --- | --- | --- | --- |
| rs12612492 | *AC008073.7* | Hippocampus | 4.68E-08 | 0.0717 | 0.893 |
| Rs9586 | *AMT* | Hypothalamus | 1.57E-08 | 0.173 | 0.942 |
| rs1850344 | *RP11-115H18.1* | Putamen Basal Ganglia | 1.84E-08 | 0.0344 | 0.859 |
| rs4840157 | *RP3-467N11.1* | Cerebellum | 1.02E-07 | 0.0278 | 0.829 |
| rs11764590 | *FTSJ2* | Caudate basal ganglia | 9.48E-11 | 1.3E-06 | 0.441 |
| rs6463710 | *RPA3* | Caudate basal ganglia | 1.56E-06 | 0.148 | 0.909 |
| rs2715147 | *PCLO* | Nucleus accumbens basal banglia | 5.5E-07 | 0.141 | 0.914 |
| rs7853605 | *RP11-165J3.5* | Cerebellum | 1.9E-07 | 0.499 | 0.983 |
| rs11030009 | *DNF-AS* | Frontal Cortex | 1.9E-06 | 1.000 | 1.000 |
| rs11039149 | *LRP4* | Nucleus accumbens basal ganglia | 1.88E-13 | 0.00204 | 0.824 |
| rs174549 | *FADS3* | Cerebellar Hemisphere | 0.00035 | 0.129 | 0.82 |
| rs2013515 | *RP11-74E22.6* | Caudate basal ganglia | 1.27E-10 | 1.000 | 1.000 |
| rs12945855 | *TTC19* | Nucleus accumbens basal ganglia | 6.75E-09 | 0.00767 | 0.788 |
| rs8072451 | *RP11-798G7.5* | Cerebellar Hemisphere | 2.04E-12 | 1.000 | 1.000 |
| Rs1292060 | *VMP1* | Cerebellar Hemisphere | 1.2E-07 | 0.0889 | 0.897 |

### Fine-mapping of TWAS association causally implicates several genes with miserableness

To identify causal genes, FOCUS was used to assign a posterior inclusion probability for genes at each TWAS region and for relevant tissue types. Across all panels, there were 95 hits included in the 90% credible gene set (Table 3). The gene, *C7orf50*, had the highest posterior inclusion probability (PIP) of 0.869 in the brain cortex. Additionally, the top multi-tissue TWAS hit from the FUSION results, *GPX1*, was included in the credible sets with a PIP of 0.192 in the frontal cortex. Several other genes also had high PIPs, such as *RP11-127L20.3* in the hippocampus with a PIP of 0.304, *MTCH2* in the nucleus accumbens with a PIP of 0.573, *ORC4* in the hypothalamus with a PIP of 0.524, *ATAD2B* in the cerebellar hemisphere with a PIP of 0.264, *FANCL* in the dorsolateral prefrontal cortex with a PIP of 0.446, and *CTC-467M3.3* in the frontal cortex with a PIP of 0.567.

**Table 3.**
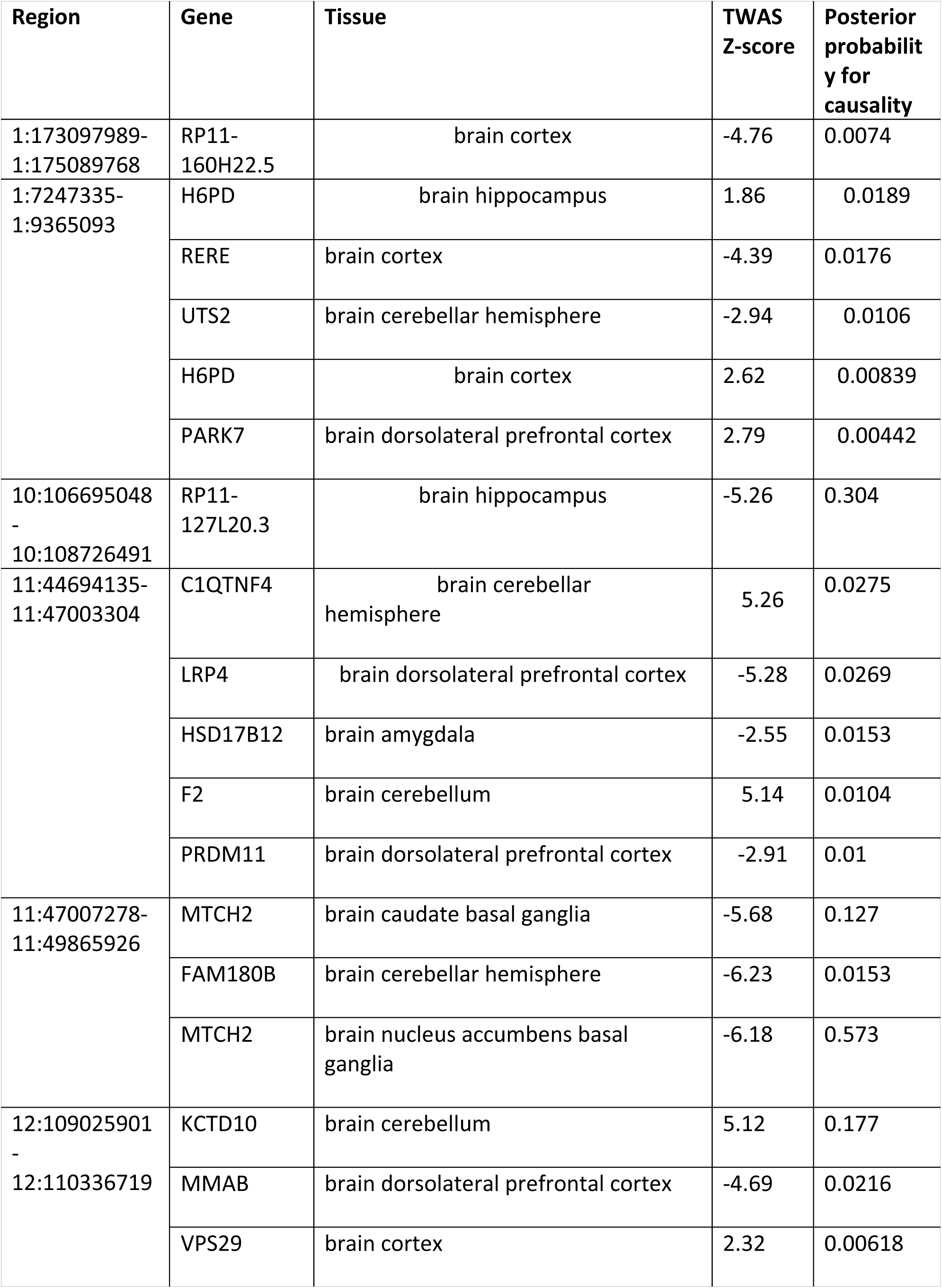
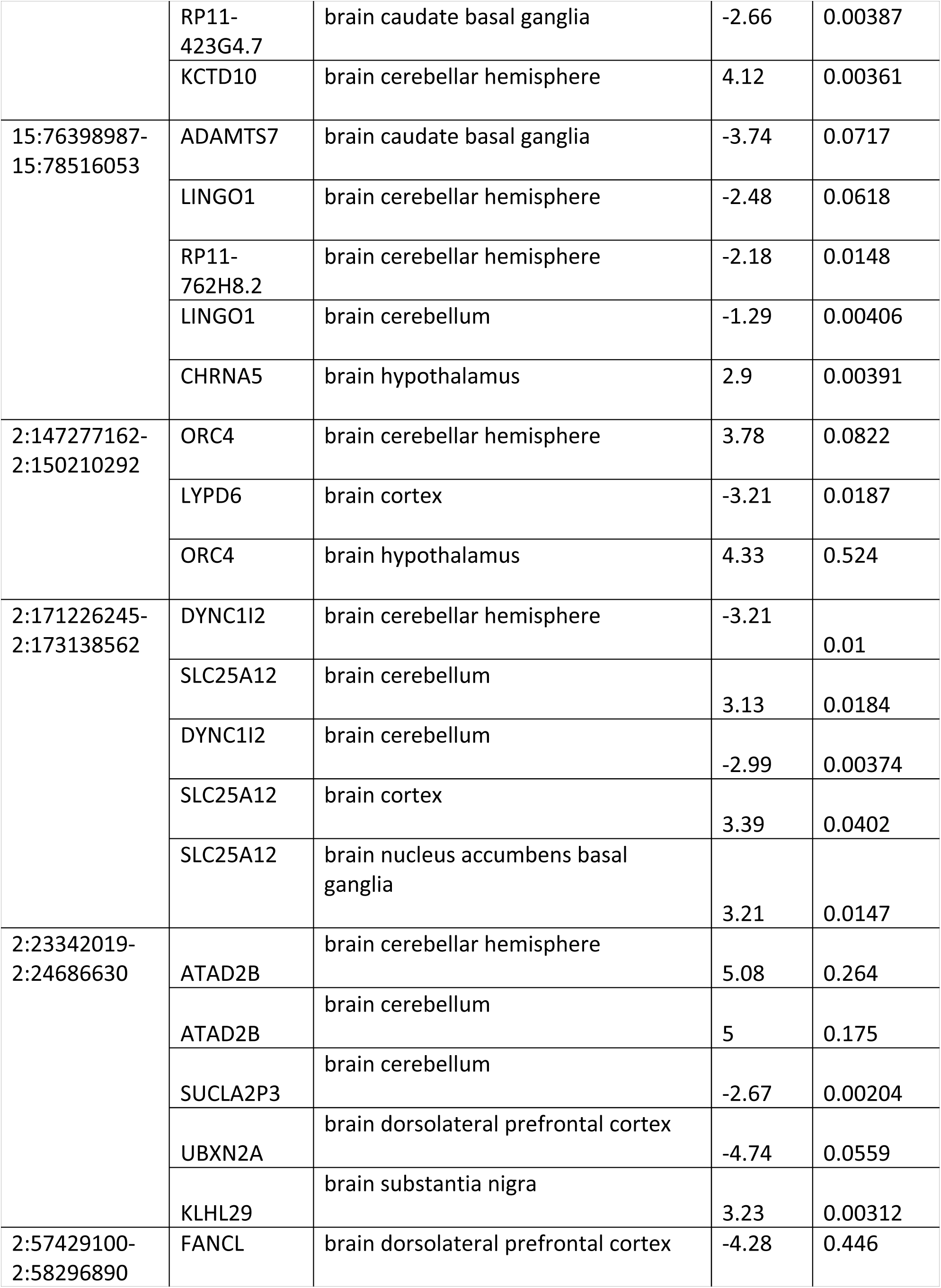
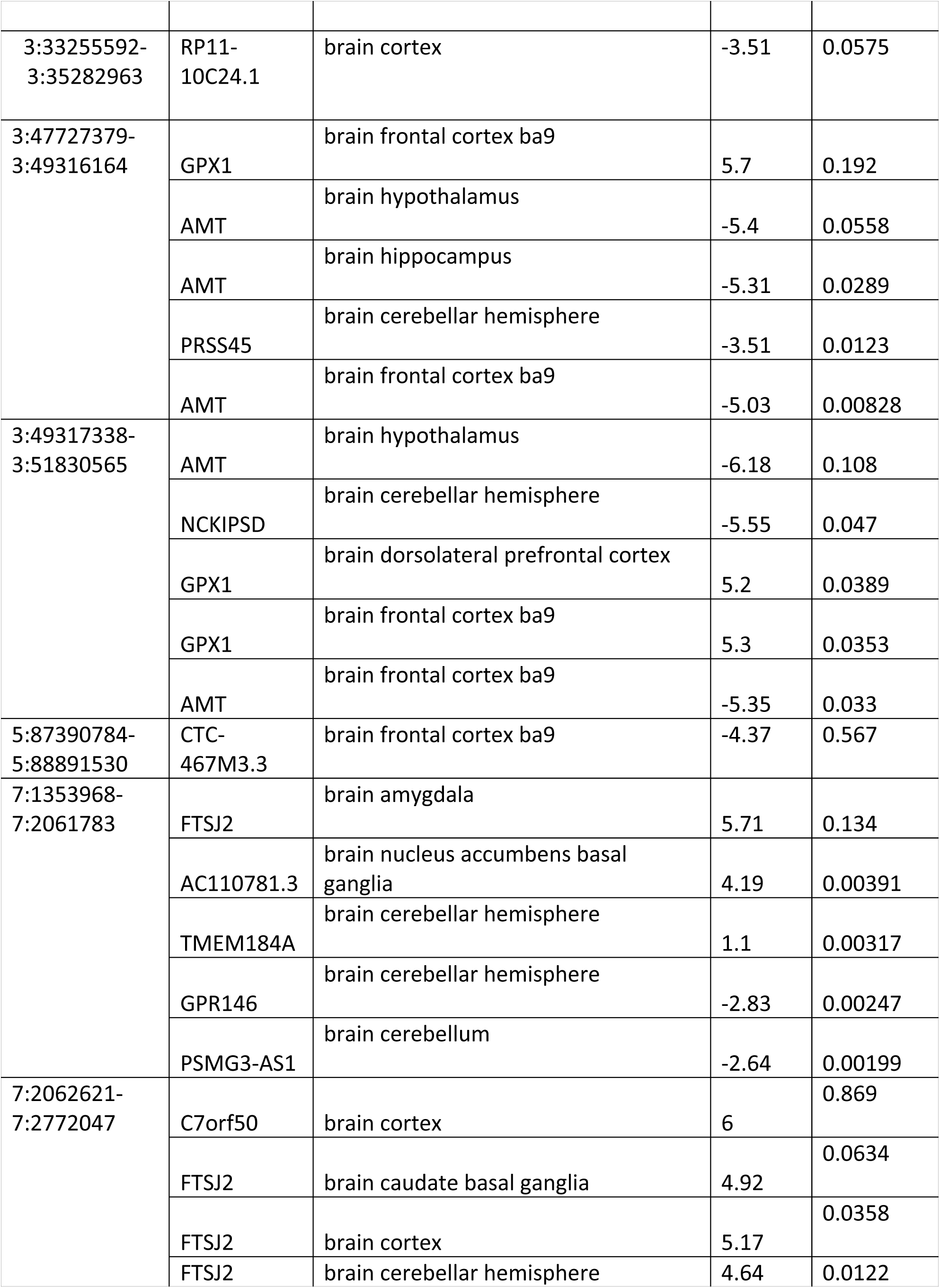
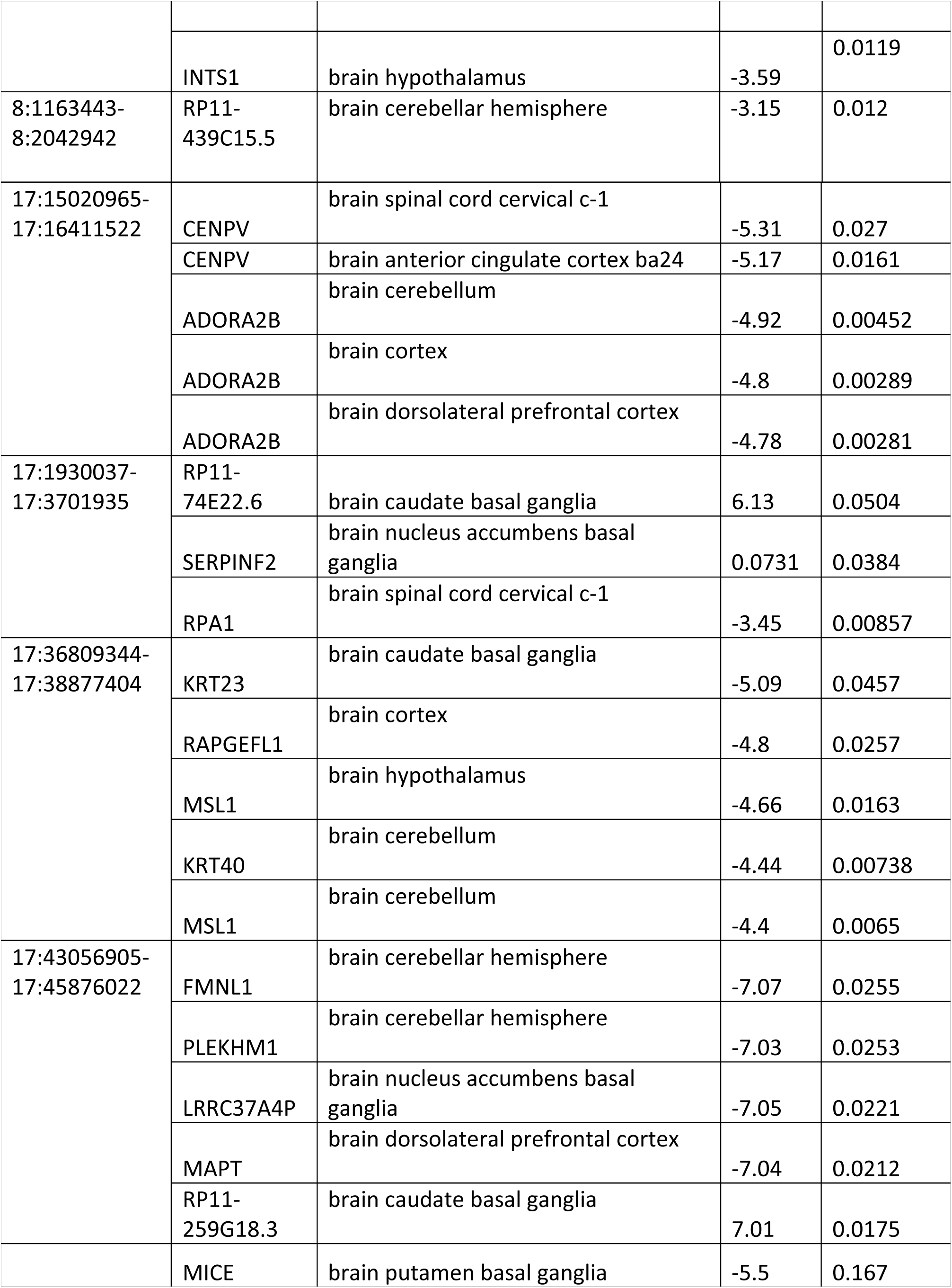
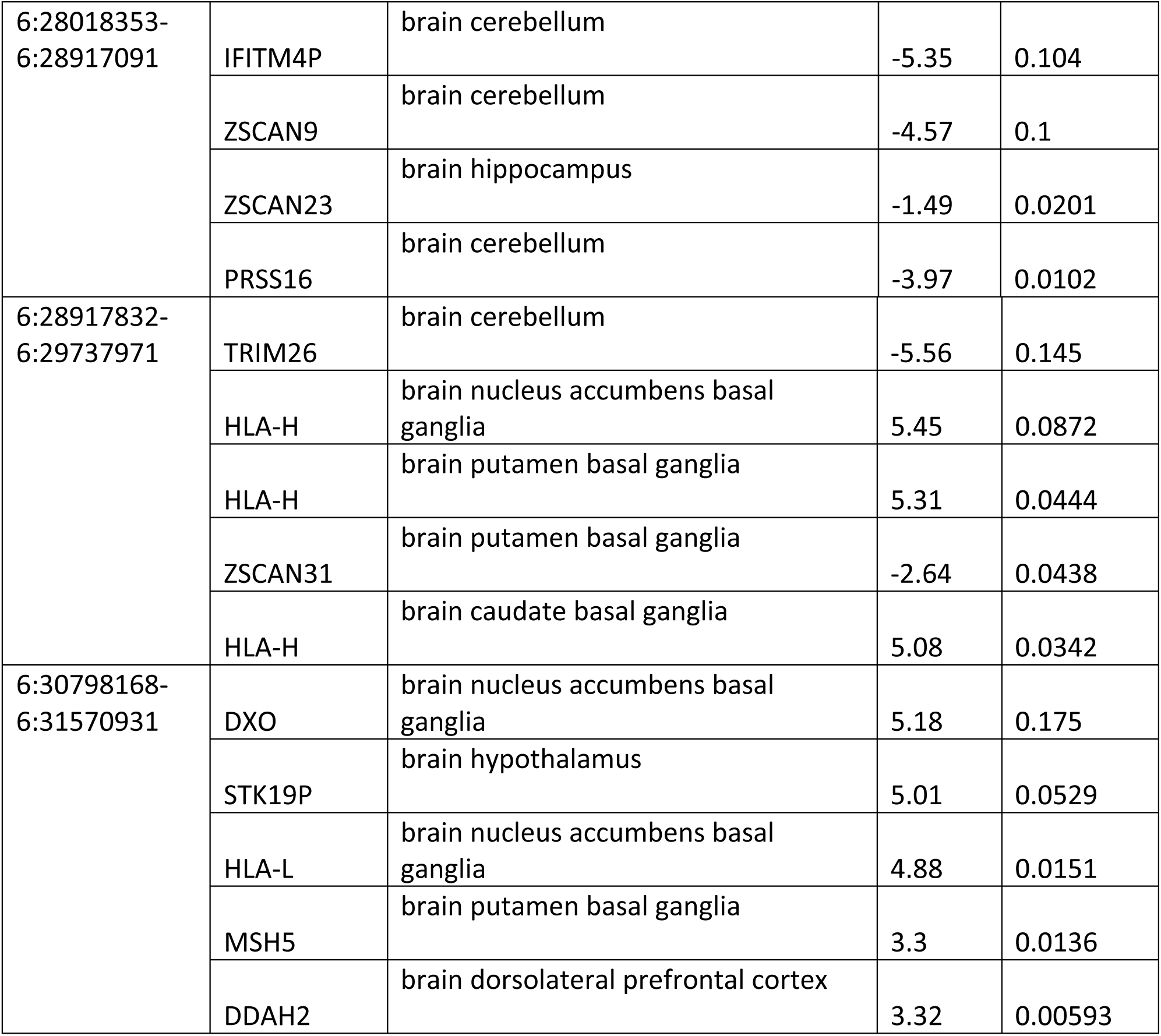
Fine-mapped genes for miserableness.

## Discussion

With the influx of several large-scale biobanks and GWAS studies, many loci are being identified. The next step is to identify biologically- and phenotypically-relevant genes found by GWAS. To date, few studies have attempted to understand the genetics of miserableness, despite the high prevalence of the trait. Here, we conducted the largest TWAS to date using the summary statistics of 373,733 individuals to further understand this neuroticism item. A total of 104 TWAS signals were transcriptome-wide significant across brain tissues with 37 unique genes.

We identified a total of 104 transcriptome-wide signals across brain tissues with 37 unique genes. The top two signals included *GPX1* frontal cortex and *RNF123* in the cerebellum. The gene, *GPX1*, encodes for a cytosolic enzyme, glutathione peroxidase-1, expressed in many different tissues^8^. This gene has previously been implicated in Alzheimer’s disease affecting cortical neurons^9^. Furthermore, neurons lacking *GPX1* leads have a greater susceptibility to oxidative-driven cell death^8^. The Z-scores for all *GPX1* hits were positive, suggesting that increased expression leads to susceptibility to miserableness. Currently, little is known about the effects of increased *GPX1* in the nervous system. The *RNF123* gene, encodes for a ubiquitin-protein ligase, and expression has been correlated with depressive disorder, which likely has genetic overlap with miserableness^10,11^.

Often, an implicated GWAS locus contains many genes. Common GWAS mapping techniques would assign the SNP association to the closest gene, however, this has been shown to be suboptimal^12–14^. After conditioning on the TWAS signal for each transcriptome-wide significant hit, most of the signal was explained by the expression of the conditioned gene.

Fine-mapping of the TWAS signals identified many genes included in the credible set, where *C7orf50* had the highest PIP of 0.869 in the brain cortex. Currently, the literature is sparse on *C7orf50*, however, querying the gene using the tissue-specific gene network (GIANT), showed potential implications in autism spectrum disorder and epilepsy^15^. For the brain cortex, GIANT implicated *CCDC85B* as the top functionally related gene to *C7orf50*. Previous studies have shown that *CCDC85B* is implicated in neural tube development^16^.

We conclude this study with some caveats and potential follow-up ideas. First, TWAS only measures the effects of cis-eQTLs and may not capture other genetic regulatory effects that contribute towards miserableness. Second, future large studies with wide ranges of phenotypes such as the All of Us initiative will allow to successfully measure the genetic susceptibility to miserableness in other ethnic populations. Here, we successfully demonstrated that behavioural traits such as miserableness have a strong genetic basis and many signals are driven by cis-eQTL. Genes such as *GPX1, RNF123, C7orf50* and *MTCH2* should be further investigated to understand the molecular consequences of dysregulated expression.

## Methods

### Genotype data and patient information

Summary statistics were obtained from Nagel et al. (2018)^4^. Details pertaining to participant ascertainment and quality control were previously reported by Nagel *et al*. (2018)^4^. Succinctly, the data was derived from the UK BioBank and miserableness was determined by asking “Do you ever feel ‘just miserable’ for no reason?”. A total of 373,733 individuals were included, with 45% of them saying “yes”. There were 47% of females who responded “yes”, and 43% of males who responded “yes”.

### Transcriptomic imputation

Transcriptomic imputation (TI) was done using eQTL panels created from tissue-specific gene expression coupled with genotypic data^17^. Here, we used all the brain tissue types from GTEx 53 v7 and the CommonMind Consortium (CMC)^6^. A strict Bonferroni-corrected study-wise threshold was used: P=4.97E-07 (0.05/100,572) (total number of genes across panels). FUSION was used to conduct the transcriptome-wide association testing^17^. The 1000 Genomes v3 LD panel was used for the TWAS. FUSION utilizes several penalized linear models such as GBLUP, LASSO, Elastic Net^17^. Furthermore, a Bayesian sparse linear mixed model (BSLMM) is used. FUSION works by computing an out-sample R^2^ to determine the best model by performing a fivefold cross-validating of every model. Further details can be found in the original manuscript.

### Conditionally testing GWAS signals

To determine how much GWAS signal remains after the expression association from TWAS is removed, joint and conditional testing was done for genome-wide Bonferroni-corrected TWAS signals. The joint and conditional analyses help to determine genes with independent genetic predictors associated with miserableness from genes that are simply co-expressed with a genetic predictor. Each miserableness GWAS SNP association was conditioned on the joint gene model one SNP at a time.

### Fine-mapping of TWAS associations

To address the issue of co-regulation in TWAS, we used the program FOCUS (Fine-mapping of causal gene sets) to directly model predicted expression correlations and to provide a posterior causal probability for genes in relevant tissue types^7^. FOCUS identifies genes for each TWAS signal to be part of a 90%-credible set while simultaneously controlling for pleiotropic effects of SNPs. Furthermore, the same TWAS reference panels for FUSION were used.

## Acknowledgements

This work was supported by a Canadian Institutes of Health Research Foundation Scheme grant (#332971). G.A.R. holds a Canada Research Chair in Genetics of the Nervous System and the Wilder Penfield Chair in Neurosciences. C.L. is a recipient of the Frederick Banting and Charles Best Canada Graduate Scholarship from the Canadian Institutes of Health Research (CIHR). C.L. conducted the experiments, analyses and drafted the manuscript. V.V. helped with analyses. A.D.L, D.S. helped with bioinformatics. P.A.D. and G.A.R. oversaw the analyses and helped draft the manuscript.

## Supplementary

**Supplementary Figure 1.**
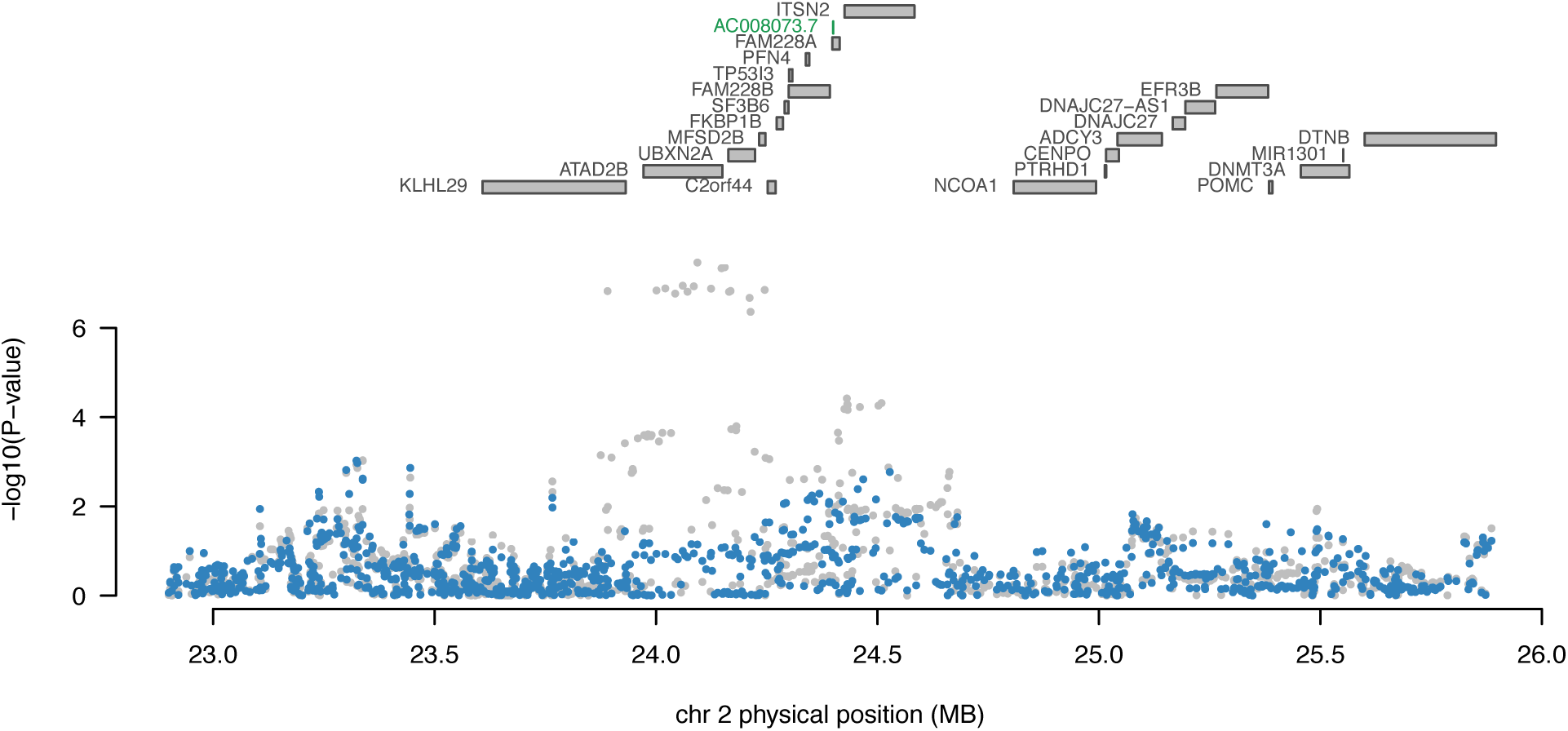
Regional association plot of chromosome 2 conditioned on *AC008073.7* expression. The top panel highlights all genes in the region. The marginally associated TWAS genes are shown in blue and the jointly significant genes are shown in green. The bottom panel shows a regional Manhattan plot of the GWAS data before (grey) and after (blue) conditioning on the predicted expression of the green genes.

**Supplementary Figure 2.**
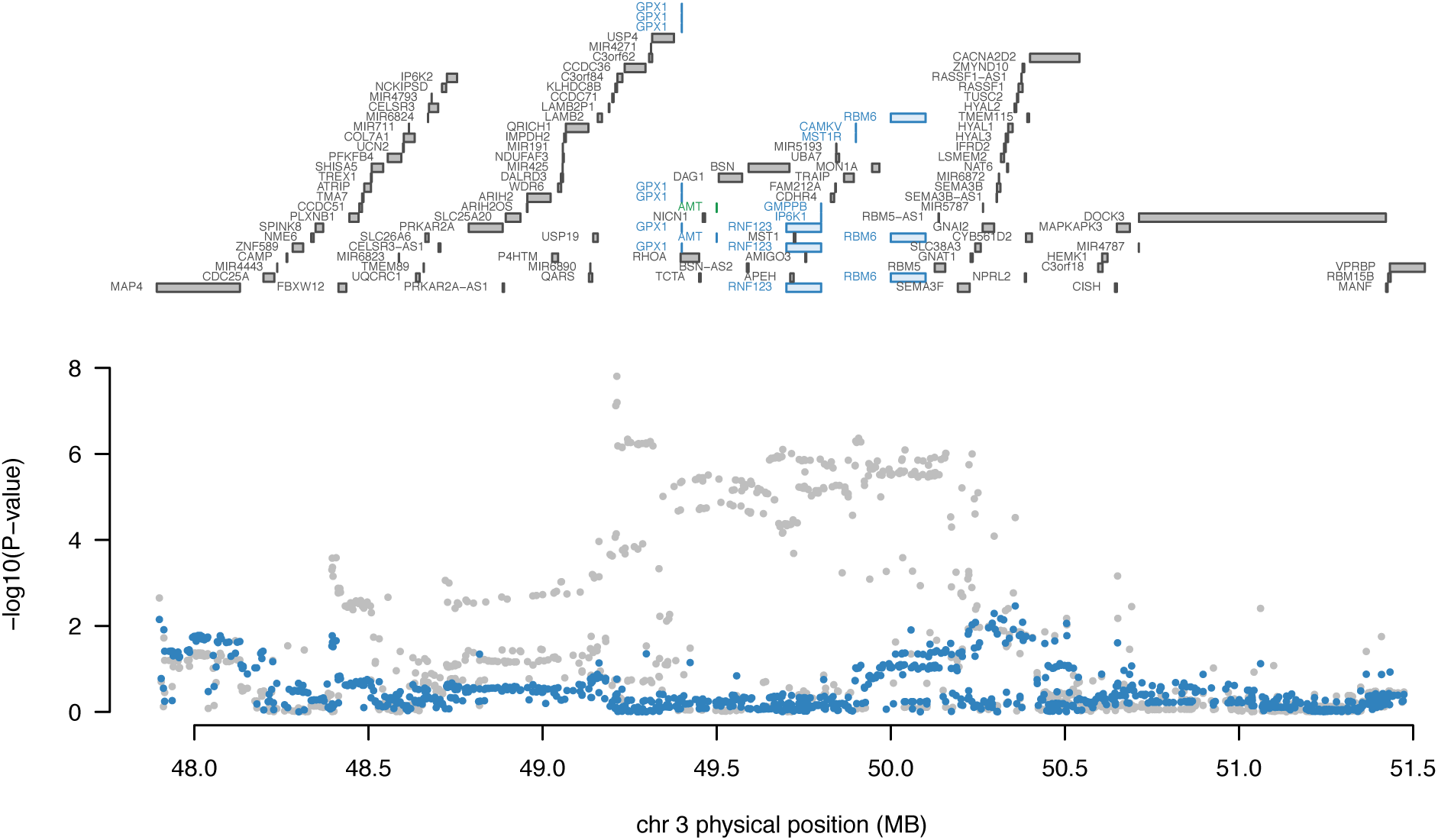
Regional association plot of chromosome 3 conditioned on *AMT* expression. The top panel highlights all genes in the region. The marginally associated TWAS genes are shown in blue and the jointly significant genes are shown in green. The bottom panel shows a regional Manhattan plot of the GWAS data before (grey) and after (blue) conditioning on the predicted expression of the green genes.

**Supplementary Figure 3.**
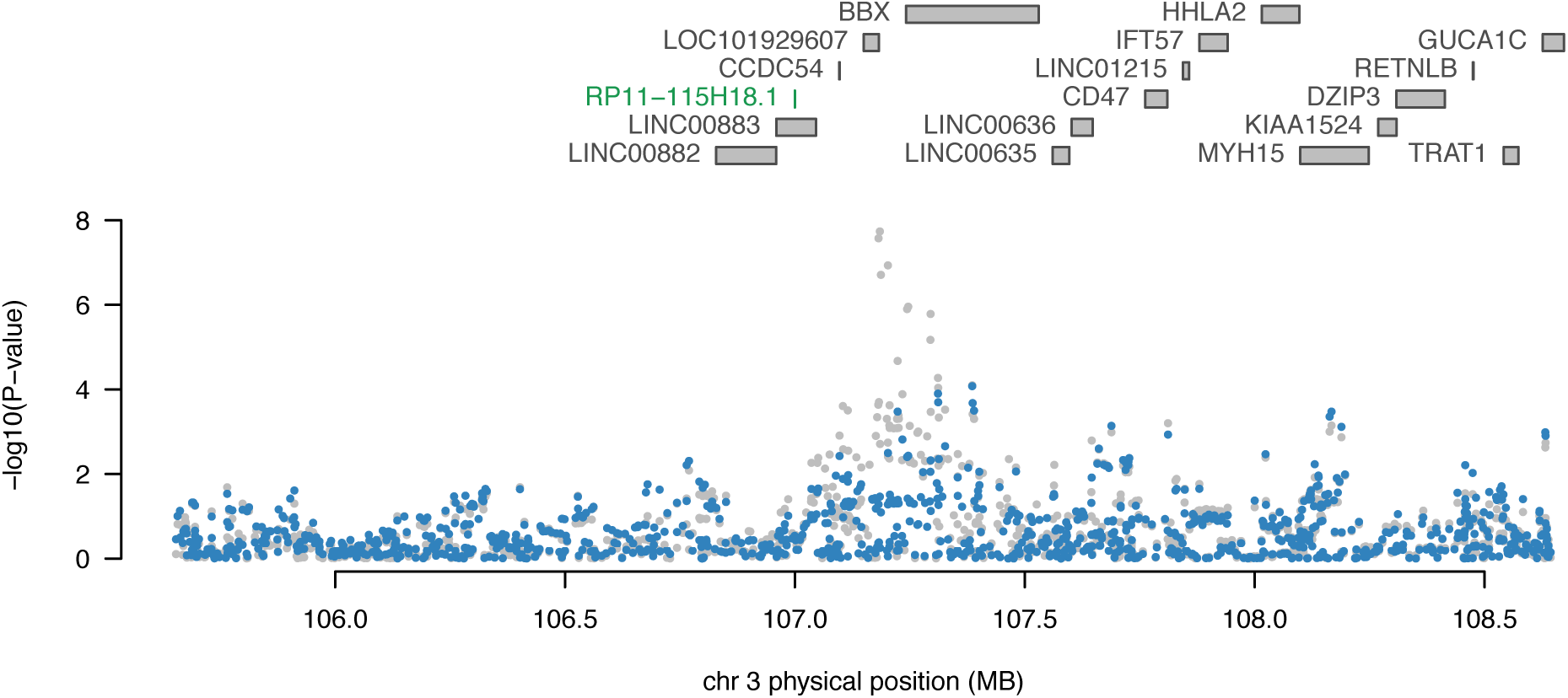
Regional association plot of chromosome 3 conditioned on *RP11-115H18.1* expression. The top panel highlights all genes in the region. The marginally associated TWAS genes are shown in blue and the jointly significant genes are shown in green. The bottom panel shows a regional Manhattan plot of the GWAS data before (grey) and after (blue) conditioning on the predicted expression of the green genes.

**Supplementary Figure 4.**
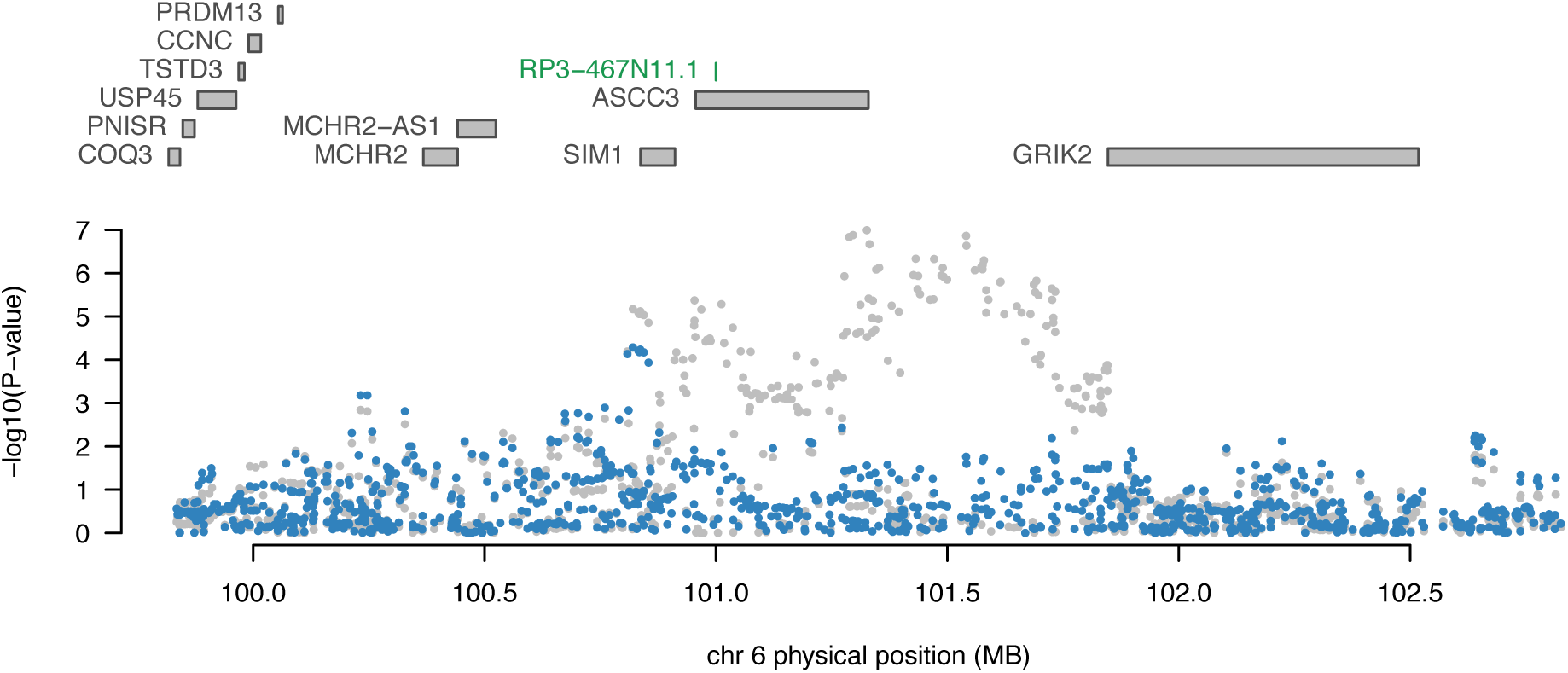
Regional association plot of chromosome 6 conditioned on *RP3-467N11.1* expression. The top panel highlights all genes in the region. The marginally associated TWAS genes are shown in blue and the jointly significant genes are shown in green. The bottom panel shows a regional Manhattan plot of the GWAS data before (grey) and after (blue) conditioning on the predicted expression of the green genes.

**Supplementary Figure 5.**
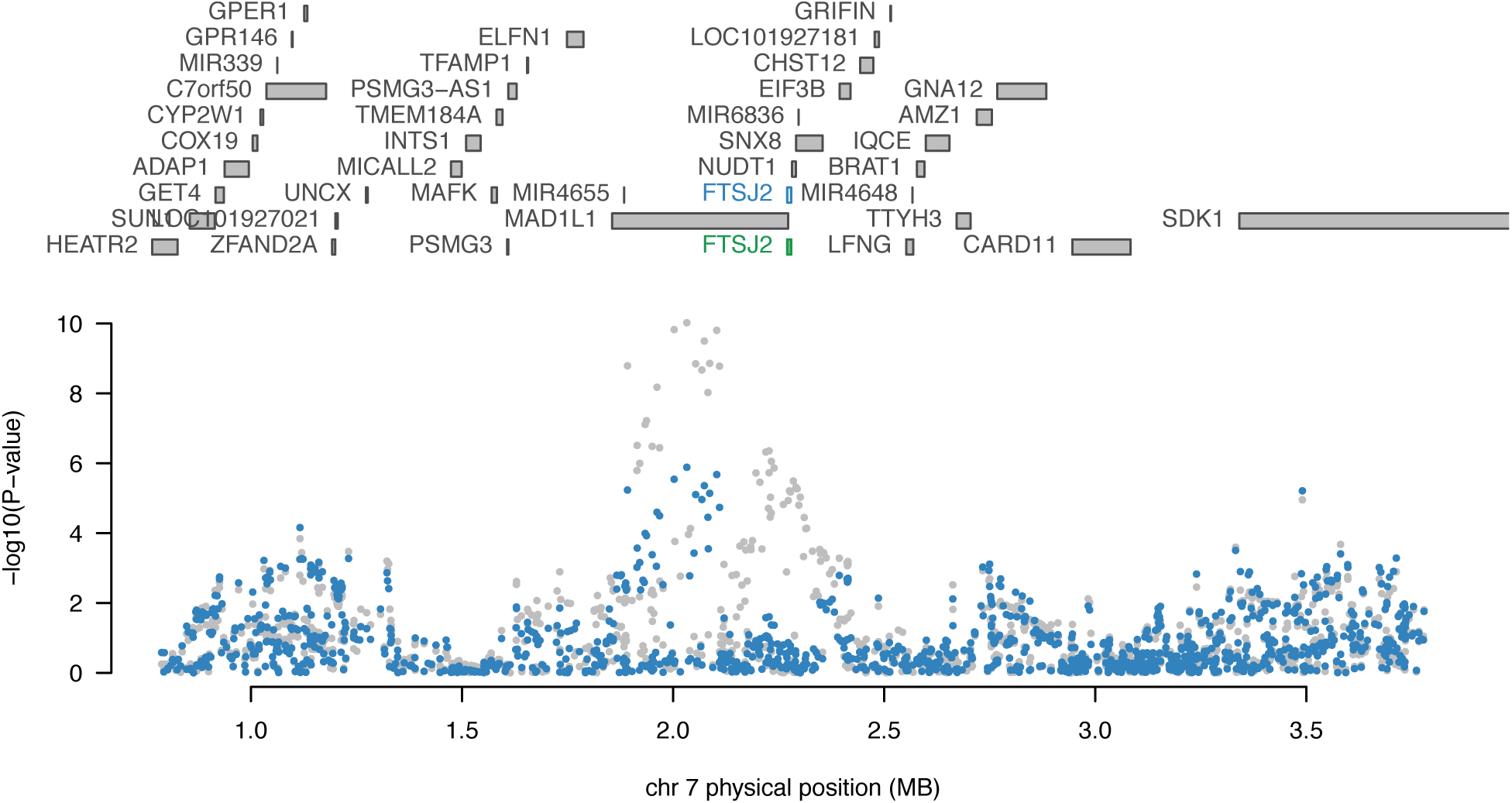
Regional association plot of chromosome 7 conditioned on *FTSJ2* expression. The top panel highlights all genes in the region. The marginally associated TWAS genes are shown in blue and the jointly significant genes are shown in green. The bottom panel shows a regional Manhattan plot of the GWAS data before (grey) and after (blue) conditioning on the predicted expression of the green genes.

**Supplementary Figure 6.**
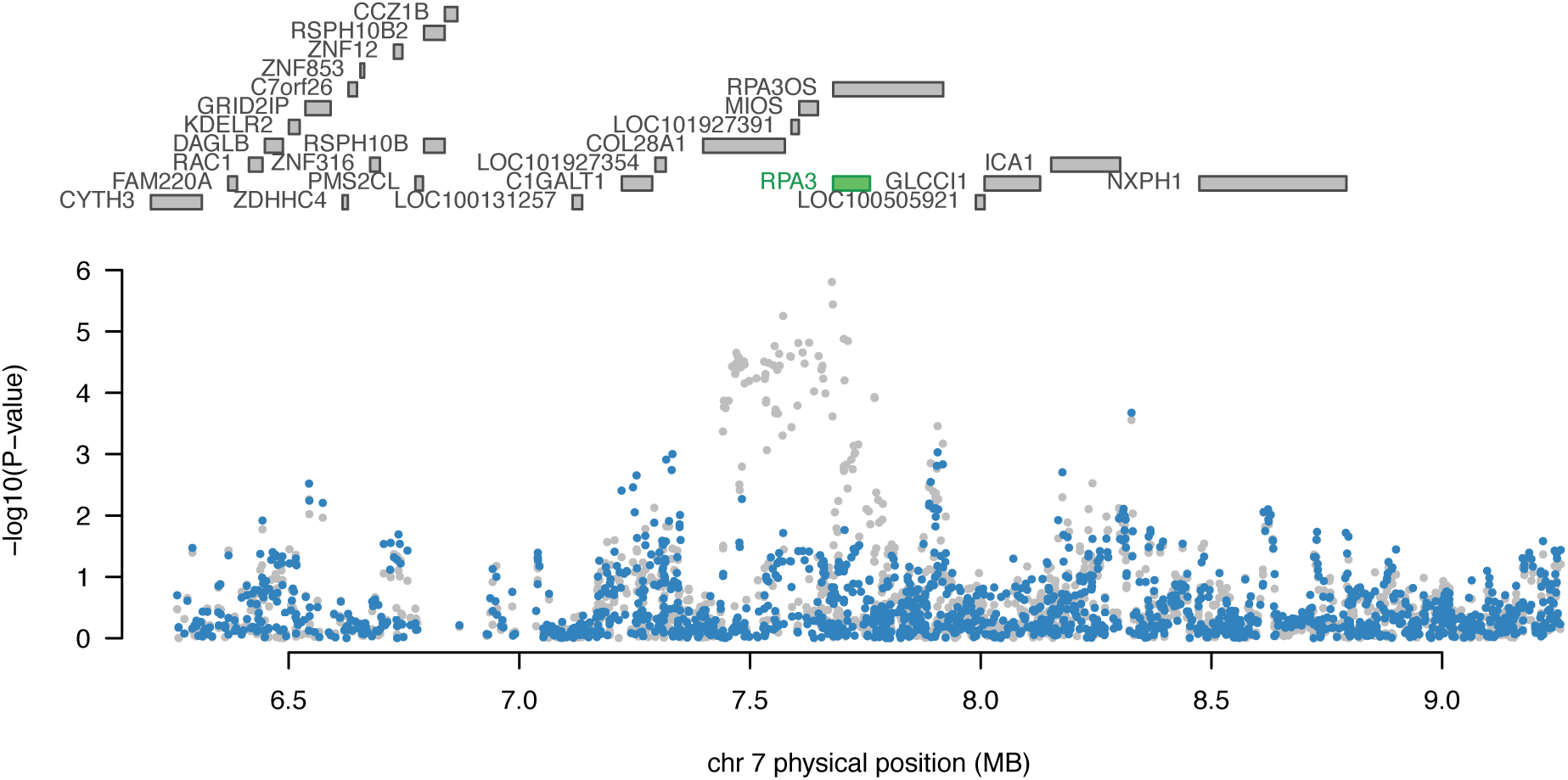
Regional association plot of chromosome 7 conditioned on *RPA3* expression. The top panel highlights all genes in the region. The marginally associated TWAS genes are shown in blue and the jointly significant genes are shown in green. The bottom panel shows a regional Manhattan plot of the GWAS data before (grey) and after (blue) conditioning on the predicted expression of the green genes.

**Supplementary Figure 7.**
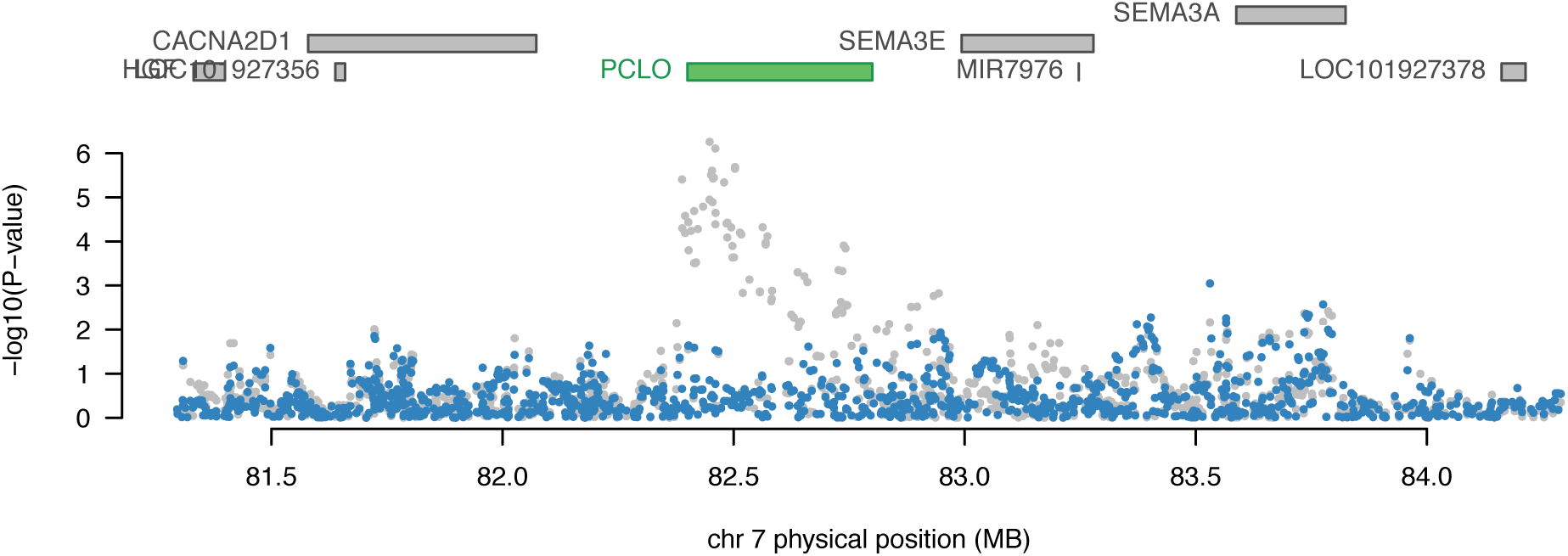
Regional association plot of chromosome 7 conditioned on *PCLO* expression. The top panel highlights all genes in the region. The marginally associated TWAS genes are shown in blue and the jointly significant genes are shown in green. The bottom panel shows a regional Manhattan plot of the GWAS data before (grey) and after (blue) conditioning on the predicted expression of the green genes.

**Supplementary Figure 8.**
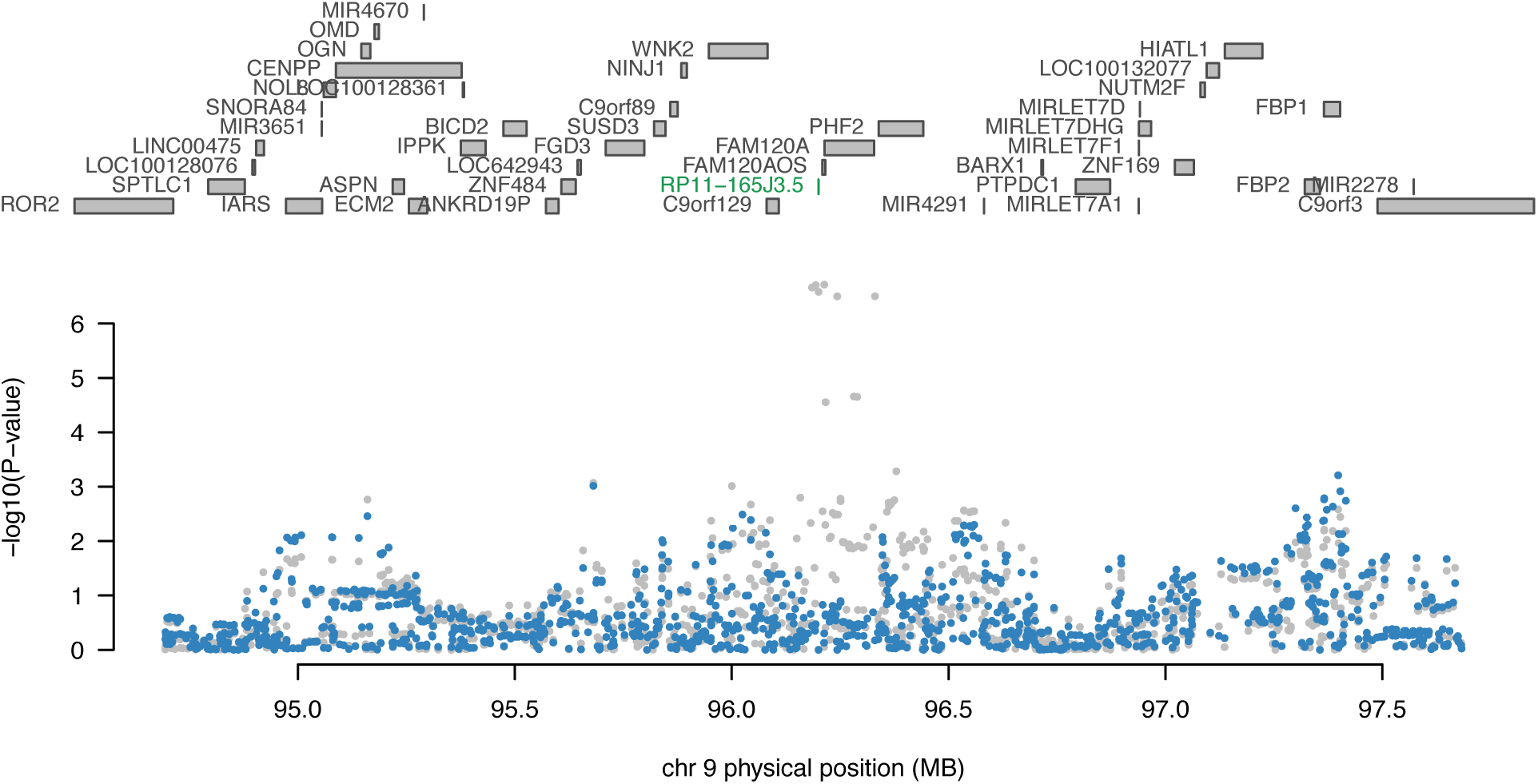
Regional association plot of chromosome 9 conditioned on *RP11-165J3.5* expression. The top panel highlights all genes in the region. The marginally associated TWAS genes are shown in blue and the jointly significant genes are shown in green. The bottom panel shows a regional Manhattan plot of the GWAS data before (grey) and after (blue) conditioning on the predicted expression of the green genes.

**Supplementary Figure 9.**
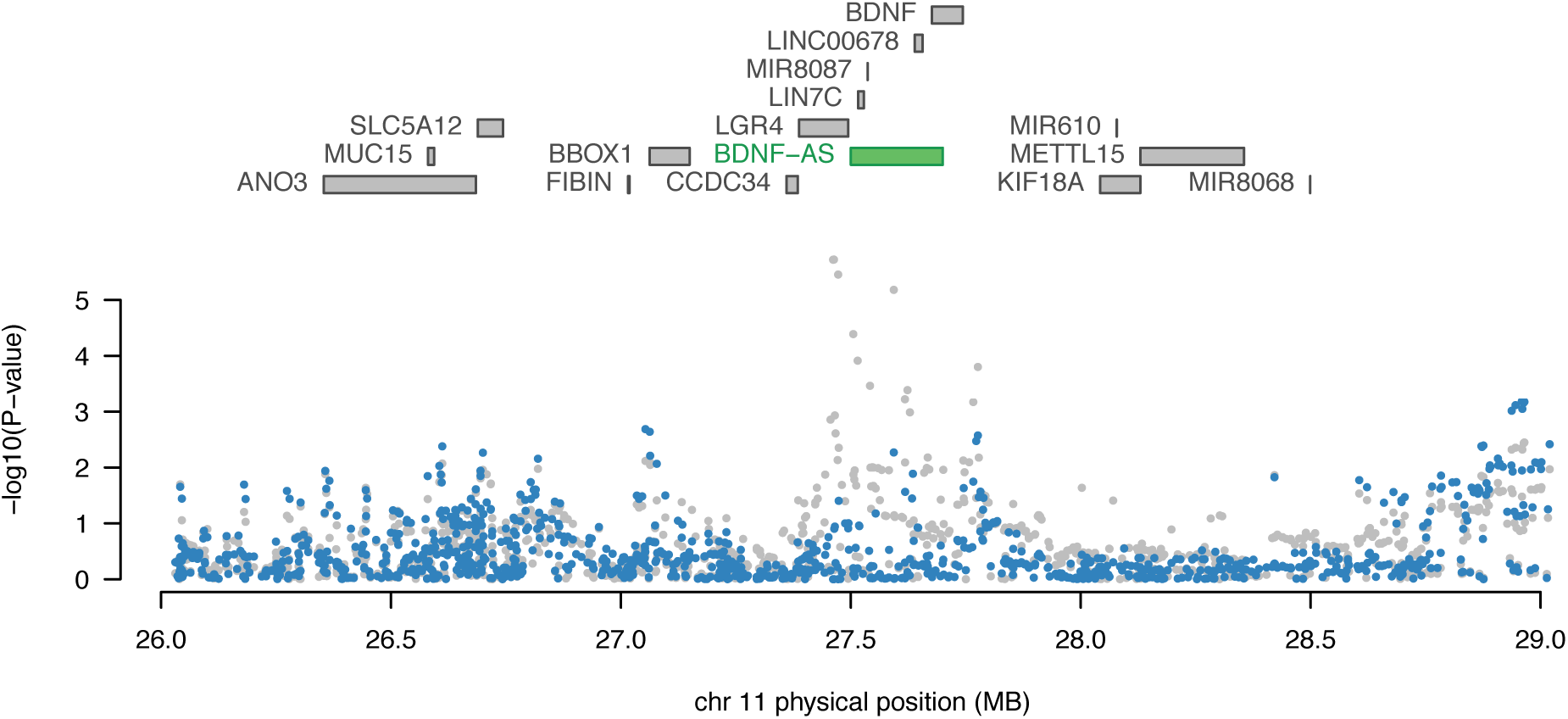
Regional association plot of chromosome 11 conditioned on *BDNF-AS* expression. The top panel highlights all genes in the region. The marginally associated TWAS genes are shown in blue and the jointly significant genes are shown in green. The bottom panel shows a regional Manhattan plot of the GWAS data before (grey) and after (blue) conditioning on the predicted expression of the green genes.

**Supplementary Figure 10.**
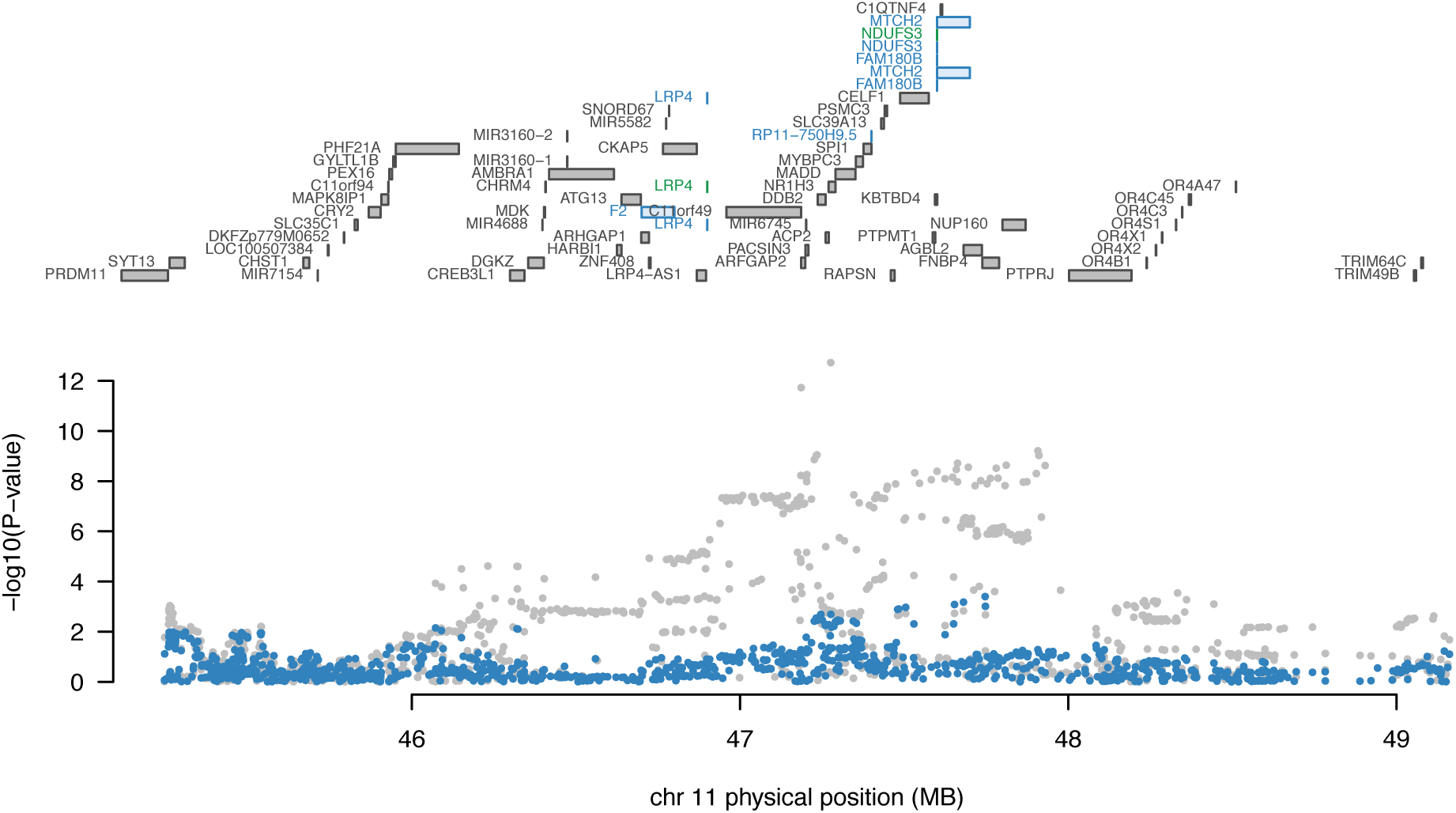
Regional association plot of chromosome 11 conditioned on *LRP4* expression. The top panel highlights all genes in the region. The marginally associated TWAS genes are shown in blue and the jointly significant genes are shown in green. The bottom panel shows a regional Manhattan plot of the GWAS data before (grey) and after (blue) conditioning on the predicted expression of the green genes.

**Supplementary Figure 11.**
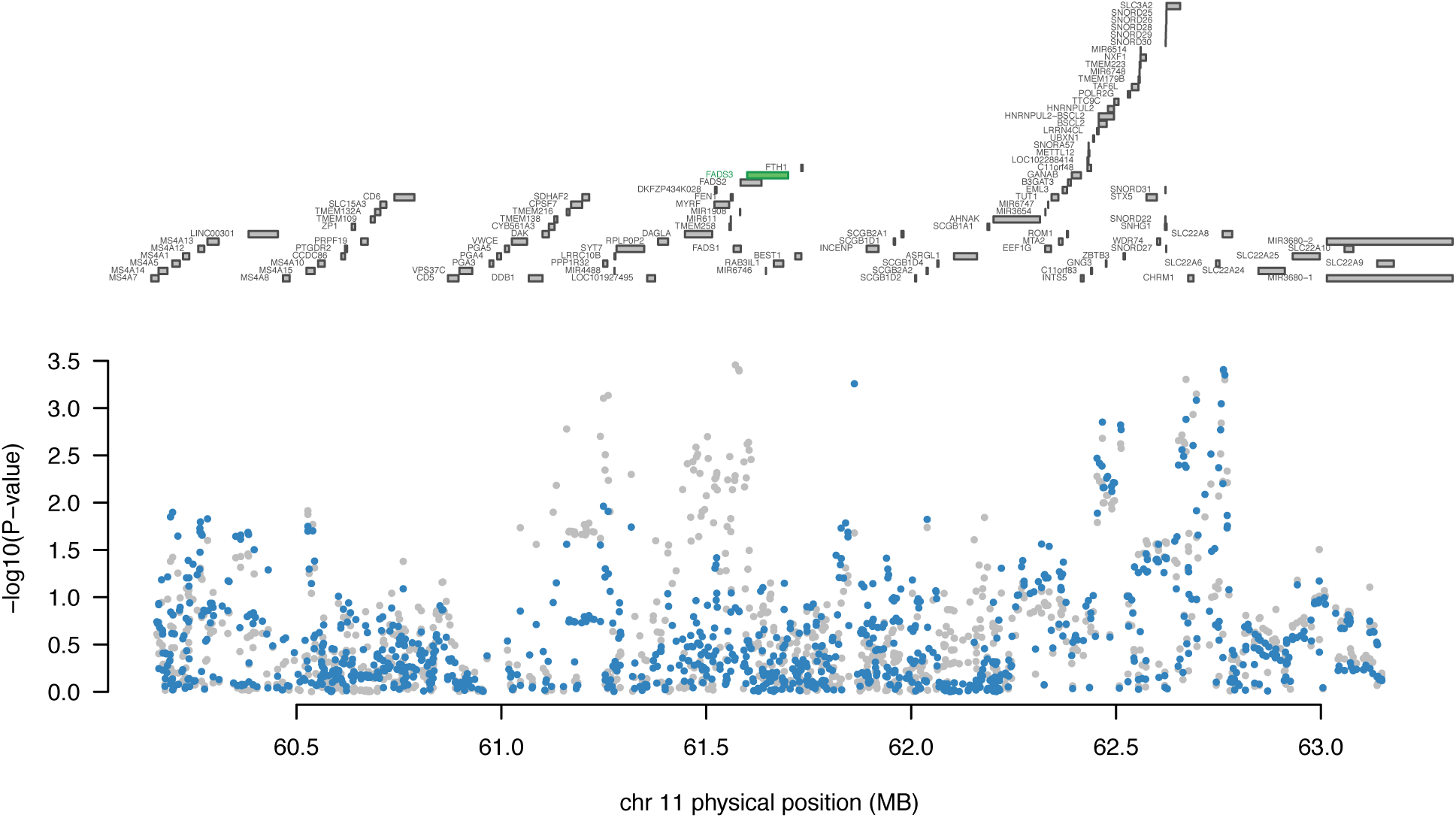
Regional association plot of chromosome 11 conditioned on *FADS3* expression. The top panel highlights all genes in the region. The marginally associated TWAS genes are shown in blue and the jointly significant genes are shown in green. The bottom panel shows a regional Manhattan plot of the GWAS data before (grey) and after (blue) conditioning on the predicted expression of the green genes.

**Supplementary Figure 12.**
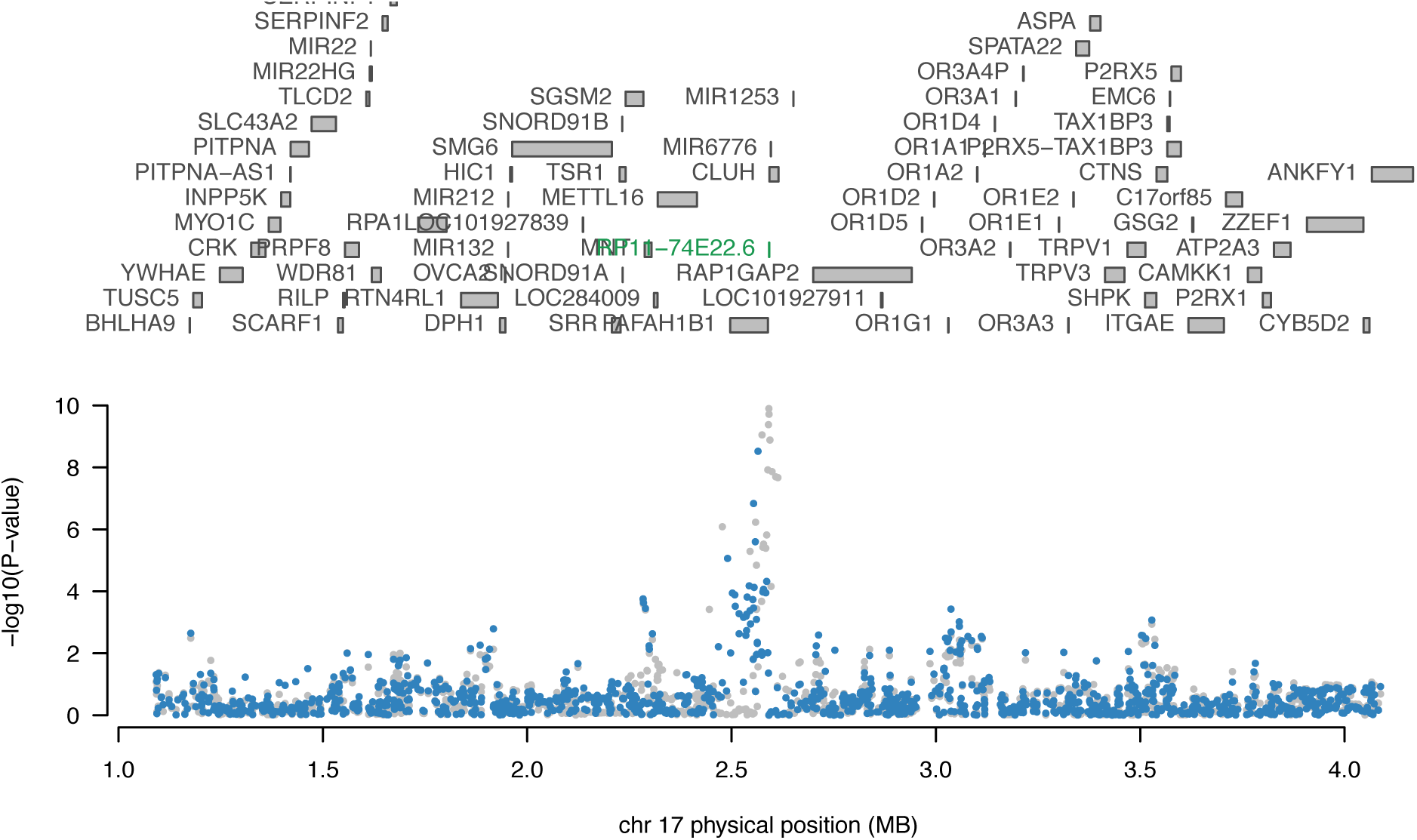
Regional association plot of chromosome 17 conditioned on *RP11-74E22.6* expression. The top panel highlights all genes in the region. The marginally associated TWAS genes are shown in blue and the jointly significant genes are shown in green. The bottom panel shows a regional Manhattan plot of the GWAS data before (grey) and after (blue) conditioning on the predicted expression of the green genes.

**Supplementary Figure 13.**
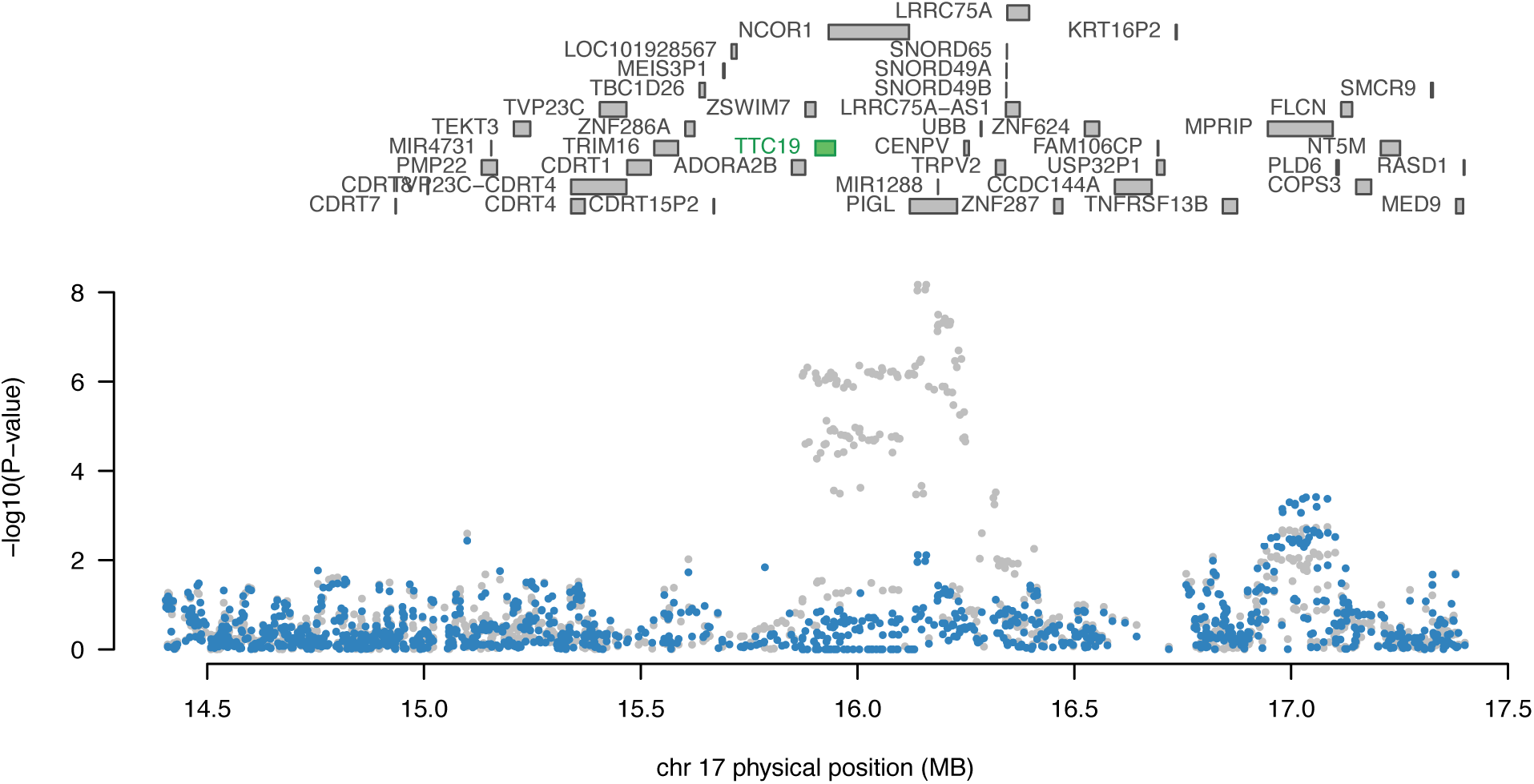
Regional association plot of chromosome 17 conditioned on *TTC19* expression. The top panel highlights all genes in the region. The marginally associated TWAS genes are shown in blue and the jointly significant genes are shown in green. The bottom panel shows a regional Manhattan plot of the GWAS data before (grey) and after (blue) conditioning on the predicted expression of the green genes.

**Supplementary Figure 14.**
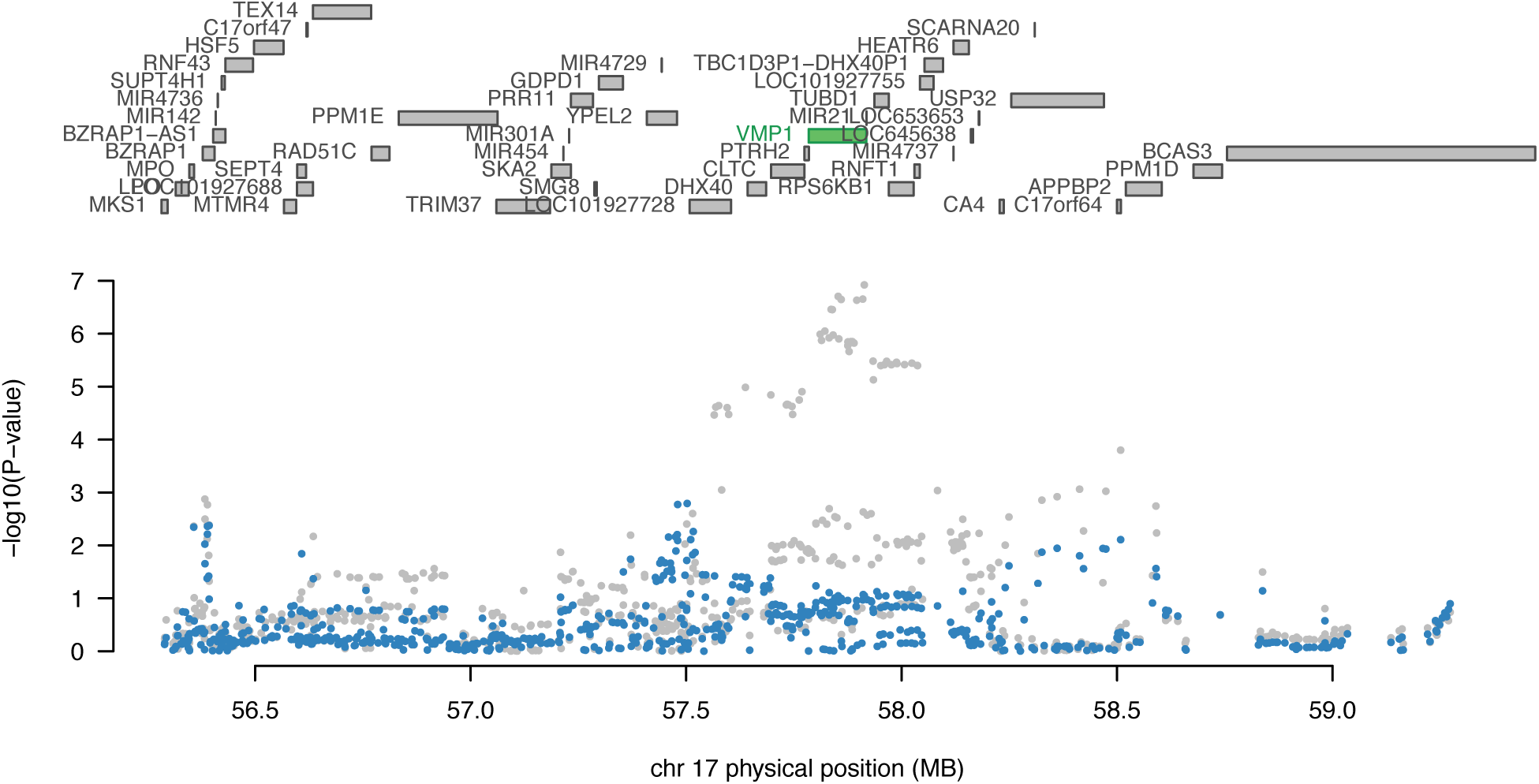
Regional association plot of chromosome 17 conditioned on *VMP1* expression. The top panel highlights all genes in the region. The marginally associated TWAS genes are shown in blue and the jointly significant genes are shown in green. The bottom panel shows a regional Manhattan plot of the GWAS data before (grey) and after (blue) conditioning on the predicted expression of the green genes.

**Supplementary Table 1.**
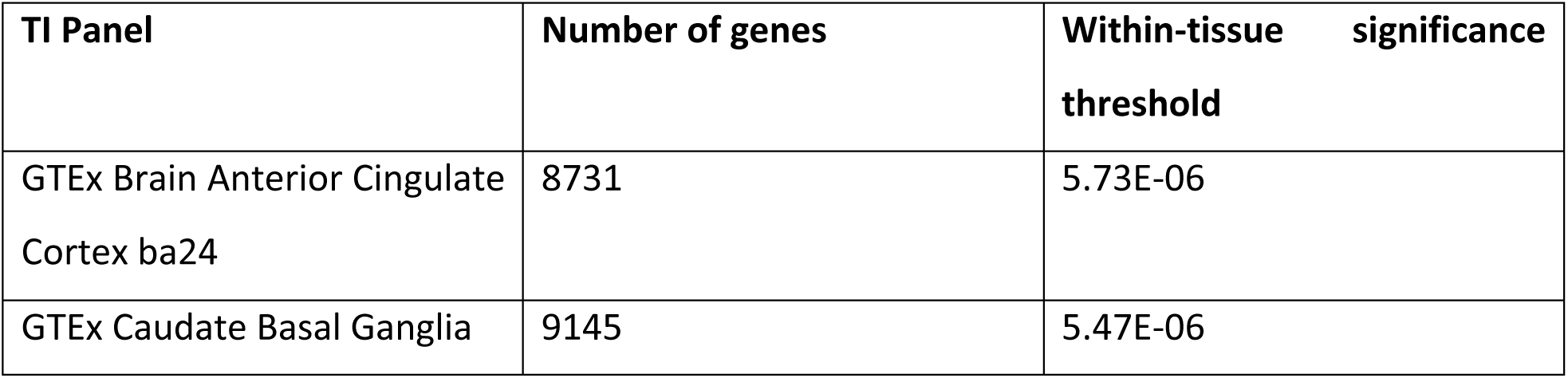
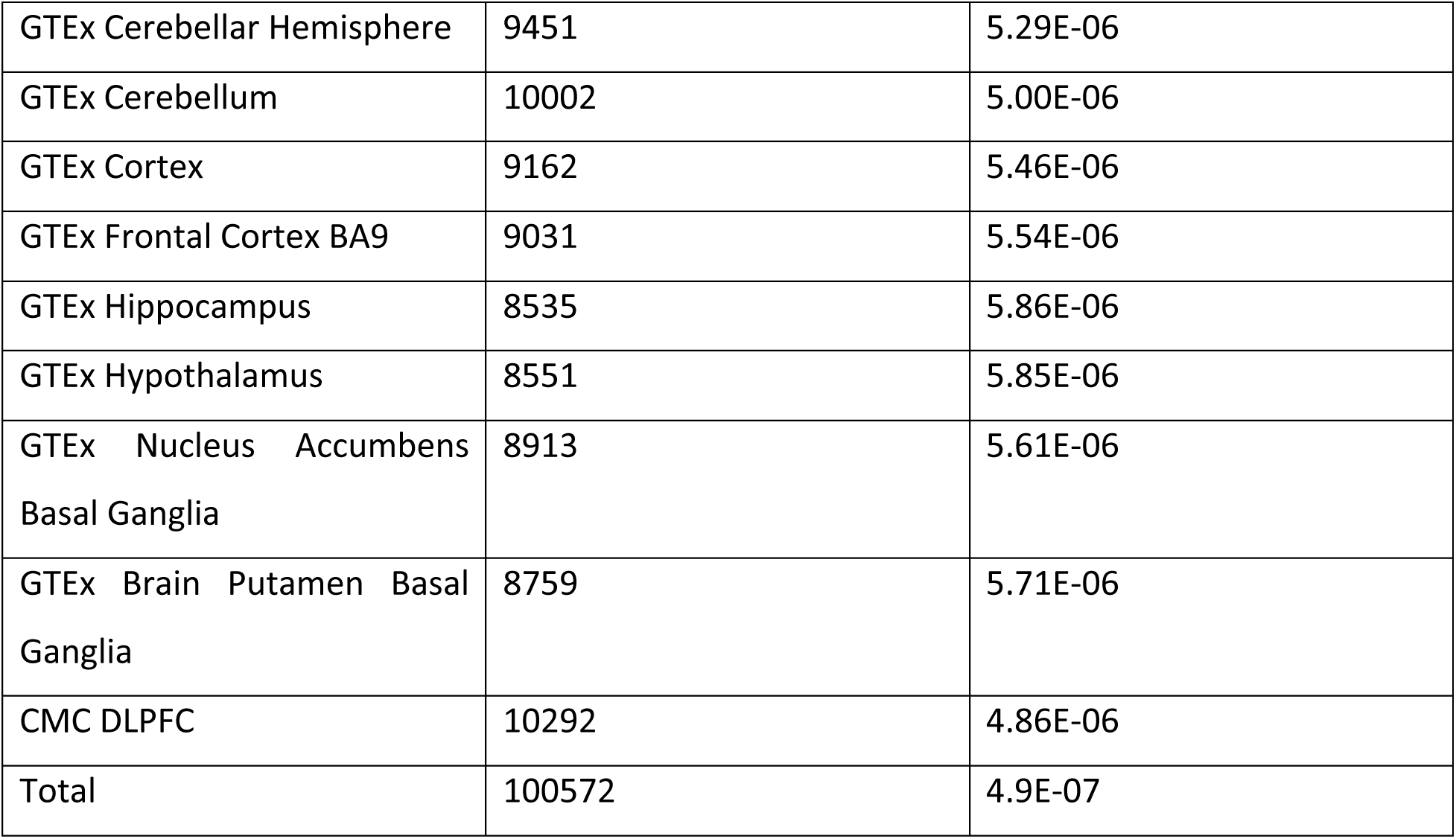
Within-tissue panel significance thresholds.

